# Tinnitus perception is linked to arousal system dysfunction

**DOI:** 10.1101/2025.07.01.662596

**Authors:** Lise Hobeika, Remy Masson, Sophie Dupont, Alain Londero, Séverine Samson

## Abstract

Tinnitus, the perception of sound in the absence of an external source, affects 14% of the population and is often associated with concentration and emotional difficulties. However, the characterization of the associated cognitive difficulties remains unclear. We hypothesize that attentional complaints are due to a dysfunction of the exogenous or endogenous orientation of attention, or of the arousal system. In this study, 200 participants (100 with chronic tinnitus and 100 matched controls) completed a battery of cognitive tasks assessing attention, alertness and executive functions, including the Attentional Network Task (ANT), Sustained Attention to Response Task (SART) with mind wandering evaluations, Stroop, and Trail Making Test. Tinnitus comorbidities, including hearing loss, sleep quality, anxiety, and hyperacusis were controlled. The results showed that individuals with tinnitus had a reduced sensitivity to alert signals, and lower sustained attention abilities, both suggesting lower levels of arousal. Mind-wandering analyses revealed fewer planning-related thoughts in the tinnitus group, suggesting higher needed cognitive resources to perform the task. Contrary to prior findings, we found no evidence of deficits in executive functioning specific to tinnitus; rather, executive impairments were associated with hearing loss and sleep disturbances. Overall, these findings support the hypothesis that tinnitus is linked to a dysfunction in the arousal system—likely involving the locus coeruleus–noradrenergic network. This work proposes a new theoretical framework implicating arousal dysregulation as a core mechanism in tinnitus-related cognitive complaints.

**Significance Statement:** Tinnitus, the perception of sound without an external source, affects millions worldwide and is often accompanied by concentration difficulties. By rigorously controlling for hearing loss, sleep deprivation, and anxiety, we isolate the core cognitive changes linked to tinnitus itself. Our study reveals that attentional deficits in tinnitus primarily arise from dysregulation in the brain’s arousal system. We propose a novel integrative framework in which arousal system dysfunction underlies the cognitive and emotional symptoms associated with tinnitus, providing a new perspective for understanding the complex interactions between attention, sleep, and anxiety in this condition.

## Introduction

Continuous non pulsatile tinnitus i.e. Subjective Tinnitus, often referred to as ‘ringing in the ears’, is a symptom characterized by the perception of sound in the absence of an external stimulus or internal sound source. This intriguing symptom is prevalent, affecting approximately 14% of the adult population (1). While tinnitus is not bothersome for most individuals, it can be highly distressing for some experiencing psychosocial disturbances such as anxiety, depression, and sleep disorders (2). Many individuals also report cognitive difficulties, particularly with concentration, describing an inability to filter this phantom perception out of their attentional focus (3). These concentration difficulties suggest possible disruptions in attentional processes, which may play a key role in the cognitive and emotional burden associated with tinnitus (4, 5).

To better understand these disruptions, it is useful to consider the attentional system as composed of multiple interconnected subsystems (6). Exogenous attention involves the involuntary capture of attention by unexpected or salient events, while endogenous attention refers to the voluntary allocation of cognitive resources toward goal-relevant stimuli. Both types of attention are conditioned by individuals’ arousal state, mediating sustained attention (tonic arousal) and alert responses (phasic arousal) (7). Some theoretical models also include the contribution of executive control within the attentional system, involving inhibitory or flexibility mechanisms (8). In tinnitus patients, reported difficulties in concentration may reflect impairments in any of these subsystems. Consequently, a hyper-responsive exogenous attention could lead to an indiscriminate attention capture by irrelevant inputs, including the tinnitus sound. A dysfunction of the endogenous system could limit the capacity to disengage from the tinnitus percept and reorient toward relevant tasks. Alternatively, the perception of tinnitus could also disrupt the arousal stage, inducing deficits in sustained attention and/or alertness.

The literature on tinnitus and cognition has primarily focused on attention and executive functions. Some studies report attentional impairments using divided (9, 10) or selective attention tasks (11), while others describe deficits in executive processes such as inhibition and cognitive flexibility (12–14), suggesting mixed findings. As highlighted by reviews and meta-analyses (15–17), the overall strength of the evidence remains limited because of methodological weaknesses related to the variability in study protocols, the small sample sizes, and the heterogeneous participant characteristics. More critically, the lack of control for comorbidities commonly associated with tinnitus - such as sleep deprivation, depression, and anxiety - renders the interpretation of many studies limited. Even hearing loss, the primary risk factor for tinnitus perception (18), is often inadequately controlled despite its well-documented impact on cognitive functioning (19). As a result, the question of whether tinnitus is associated with cognitive dysfunction—particularly in the domains of attention and executive function—remains open.

The objective of this study was to characterize attention and executive function in individuals with chronic tinnitus (over 3 months). To this end, 200 participants with and without tinnitus completed a neuropsychological assessment including attentional and executive function tasks. The Attentional Network Task (ANT, Figure 1) (8) was used to assess endogenous and exogenous attention, the tonic and phasic arousal as well as flexibility. The Sustained Attention to Response Task (SART, Figure 2) (20) provided a precise measure of tonic arousal and a mind-wandering evaluation. The Trail Making Test (shifting) and the Stroop test (inhibition) were used to evaluate executive functions. To rigorously control for tinnitus-related comorbidities, we conducted a comprehensive assessment of hearing health (audiometry with high frequencies, otoacoustic emissions, hyperacusis, as well as tinnitus severity for persons with tinnitus), emotional functioning (anxiety and depression), stress and sleep deprivation.

**Figure 1.**
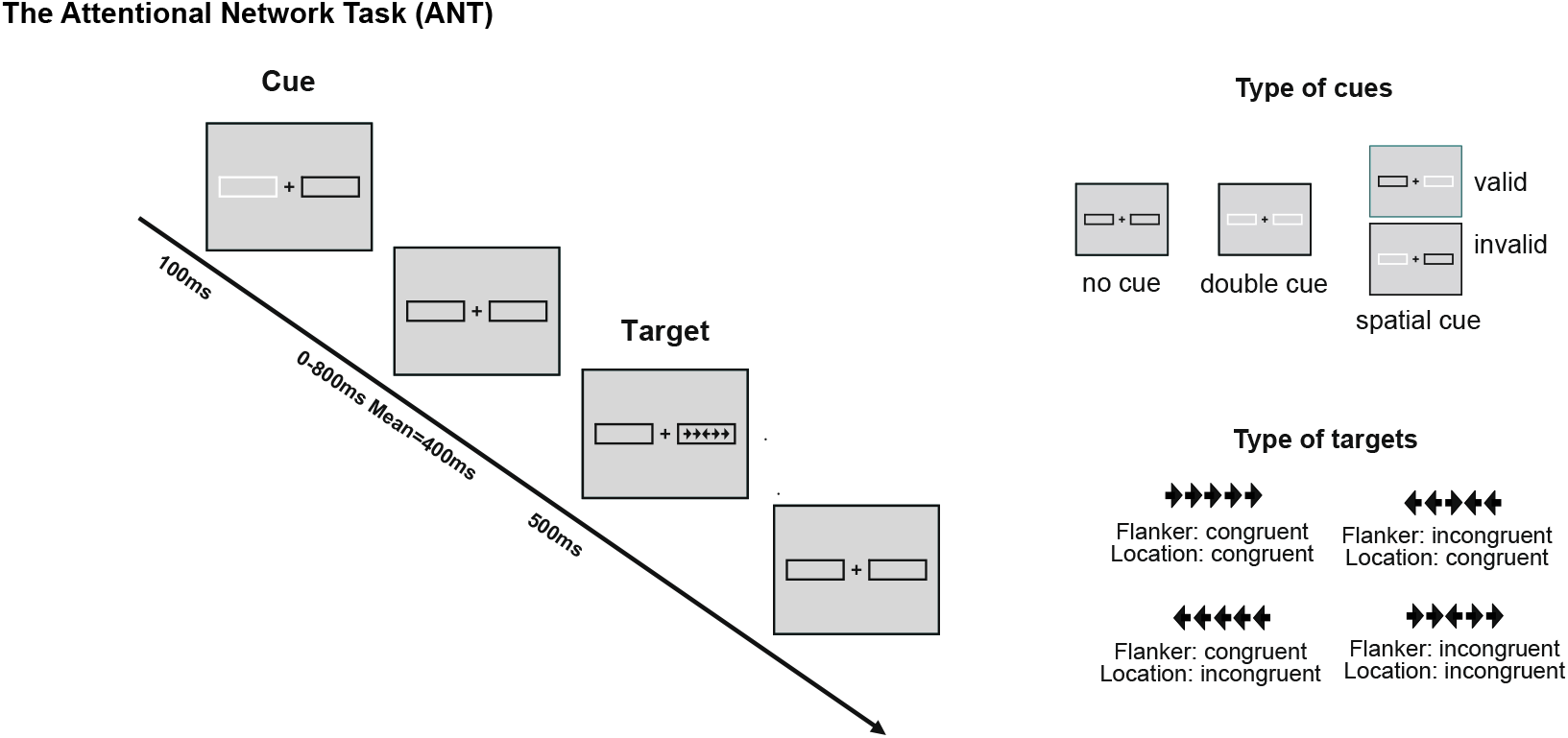
Illustration of the revised Attentional Network Test. Each trial can begin with a cue presentation, which varies depending on the type of condition (none, double, valid, or invalid). When a cue is present, a cue box flashes for 100 ms. After a variable interval (0, 400, or 800 ms), the target stimulus appears - a central arrow flanked by two arrows on each side, which can be either congruent (pointing in the same direction as the target) or incongruent (pointing in the opposite direction). The target and flankers remain on screen for 500 ms. Participants respond by indicating the direction of the central arrow. Following the response, a post-target fixation period occurs, lasting between 2000 and 12,000 ms before the next trial begins.

**Figure 2:**
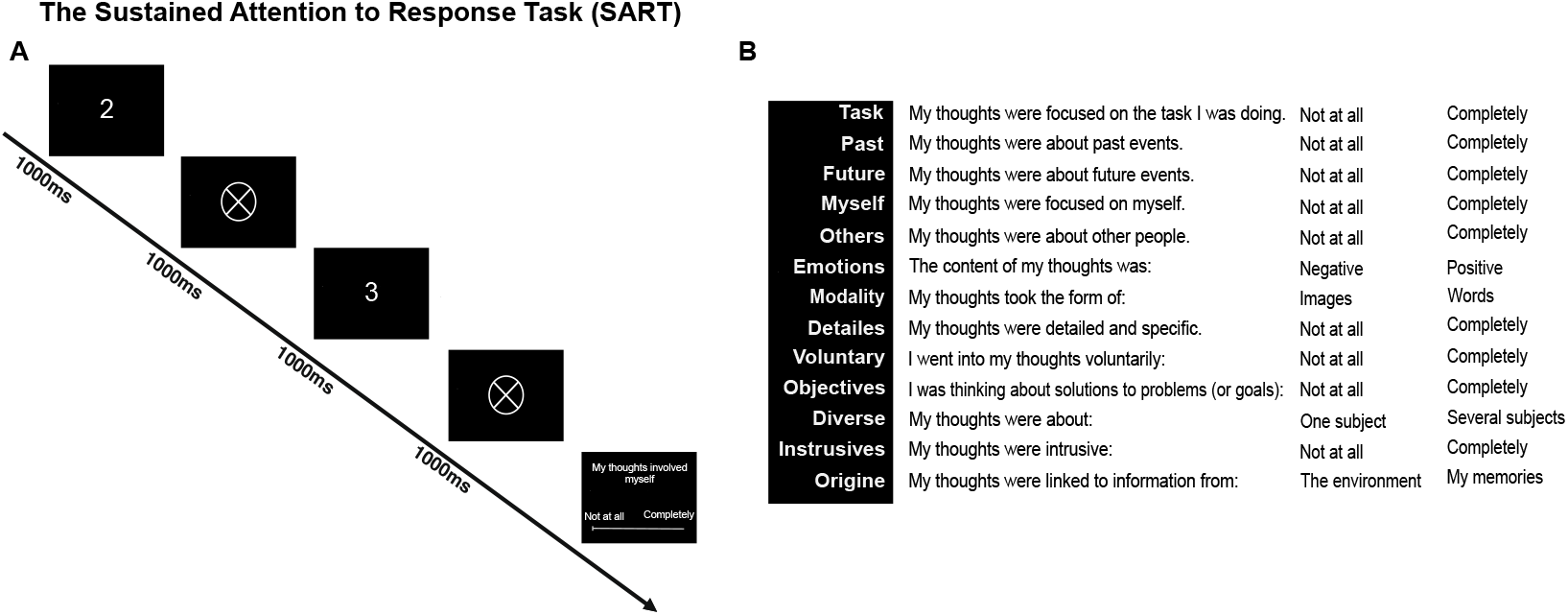
Illustration of the Sustained Attentional to Response Task. **A**. The task consisted of sequential digit presentations, ranging from 1 to 9. Participants were instructed to press a response button for each digit presentation, except when the No-Go target “3” appeared. **B**. The task was periodically interrupted by a series of 13 probe questions, randomly distributed throughout the session, which assessed the content of participants’ thoughts at the moment of interruption.

## Results

### Creation of scores capturing the emotional and hearing comorbidities

Participants with and without tinnitus did not significantly differ in terms of age, sex ratio, laterality, and level of education (Table 1). We evaluated the potential group differences in emotional functioning. Tinnitus group reported significantly higher levels of anxiety, depression, perceived stress, sleep disturbance, and hyperacusis. To control for the potential confounding effects of these comorbidities, we conducted a correlation-based data reduction analysis to identify one or several scores to include in our linear model analysis cognitive functioning, while preventing predictors multicollinearity. Among the mental health scores, we selected the STAI scores as they were highly correlated with the HADS-A, HADS-D, and PSS4 scores (r > .70, p < .001, Pearson correlation). The scores of the STAI and sleep disorders, as well as STAI and hyperacusis (Khalfa questionnaire), were correlated, but under threshold which allowed the inclusion of both scores in a linear model (r = .50, p < .001, r = .40, p < .001, respectively). Therefore, the anxiety scores (STAI), sleep disorders and hyperacusis were included as covariates in the analyses of the cognitive tasks.

**Table 1:** Tinnitus and control group descriptive data. Demographic status, emotional state, sleep disorder and hearing functioning of the two groups of participants.

|  | Control<br>N= 100 | Tinnitus<br>N= 100 | Test | p value |
| --- | --- | --- | --- | --- |
| **Demographic** |  |  |  |  |
| Age | 43 ± 13 | 45 ± 13 | Student's *t*-test | ns |
| Sex (F/M) | 54/46 | 57/43 | Pearson's chi<sup>2</sup> test | ns |
| Laterality<br>(Left/ Ambidextrous/ Right) | 13/2/85 | 12/0/88 | Pearson's chi<sup>2</sup> test | ns |
| Years of education | 15 ± 3 | 16 ± 3 | Student's *t*-test | ns |
| **Emotional state** |  |  |  |  |
| Anxiety (STAI) / 80 | 43 ± 10 | 50 ± 10 | Student's *t*-test | <.001 |
| Anxiety (HADS-A) /21 | 7 ± 4 | 10 ± 5 | Student's *t*-test | <.001 |
| Depression (HADS-D) /21 | 5 ± 4 | 7 ± 4 | Student's *t*-test | <.001 |
| Stress (PSS4) /16 | 6 ± 4 | 8 ± 3 | Student's *t*-test | <.001 |
| Sleep disorder (ISI) /28 | 6 ± 4 | 8 ± 4 | Student's *t*-test | <.001 |
| **Hearing** |  |  |  |  |
| Hyperacusis (Khalfa) /42 | 15 ± 9 | 24 ± 9 | Student's *t*-test | <.001 |
| Tinnitus severity (THI) /100 |  | 52 ± 24 |  |  |
| Tinnitus duration (years) |  | 8 ± 10 |  |  |
| Tinnitus central pitch (kHz) |  | 6,500 ± 2,600 |  |  |

Second, we measured participants hearing abilities with pure tone audiometry thresholds, evoked oto-acoustic emissions (EOAE) and distortion products otoacoustic emission (DPOEA).

*Pure tone audiometry* analysis revealed a significant effect of the factor Group (Wald χ^2^ = 3.9, p < .001), of the factor Frequency (Wald χ^2^ = 5,776, p < .001) as well as a significant interaction between Group and Frequency (Wald χ^2^ = 147, p < .001). Post-hoc tests indicated that the hearing thresholds of the Control group were significantly better than those of the Tinnitus group across multiple frequencies, particularly in the range of 3 kHz to 16 kHz (Bonferroni post-hoc tests: p < .01 for 3 kHz, p < .001 for frequencies between 4 kHz and 12 kHz) (see Figure 3.A).

**Figure 3:**
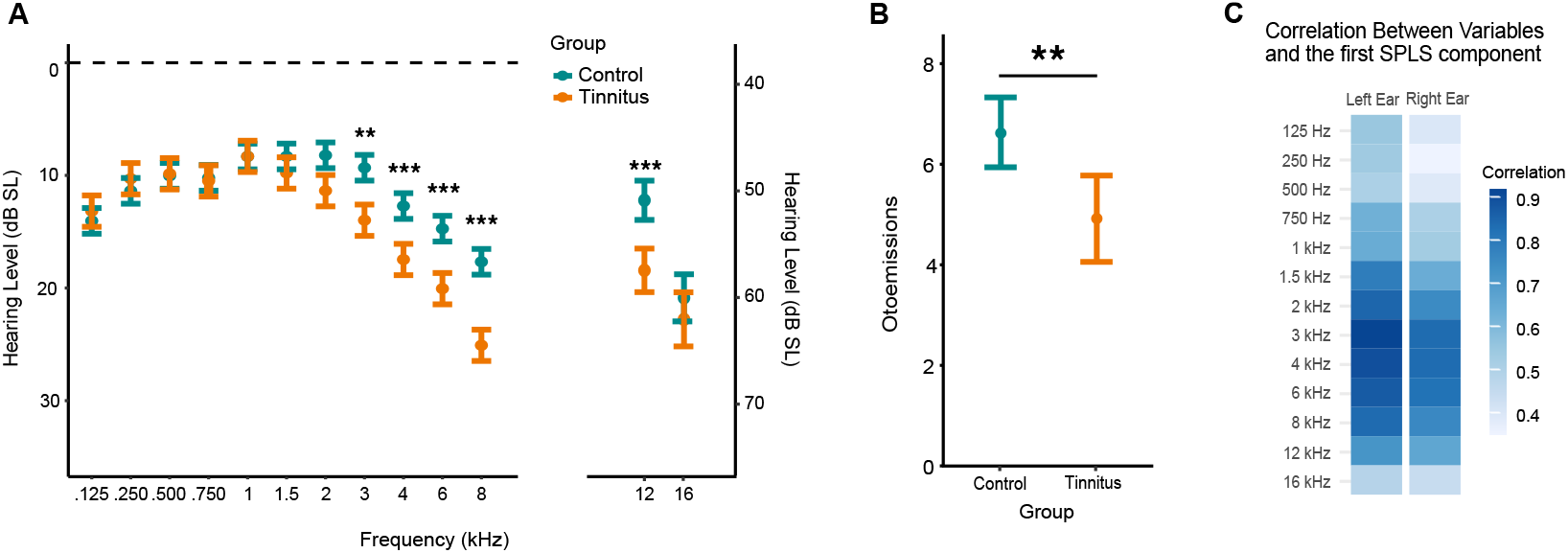
Audiological differences between tinnitus and control groups. **A**. Mean hearing thresholds (in dB SL) for tinnitus and control groups across frequencies ranging from 125Hz to 16kHz. Significant group differences were observed at frequencies from 3kHz to 12kHz. B. Mean otoacoustic emission levels showing significantly reduced responses in the tinnitus group compared to controls. C. Derivation of a composite hearing loss score using Partial Least Squares Discriminant Analysis (PLS-DA). This analysis extracted a component that optimally discriminates between control and tinnitus groups based on their hearing thresholds.

*Evoked Otoacoustic Emissions (EOAE)* analysis revealed significant effect of the tested Frequency (χ^2^(4) = 621, p < .001) with significant differences between all tested Frequencies (mean values: 6.3 ± 0.8 9.5 ± 0.8, 7.3 ± 0.8, 5.3 ± 0.8 and .5 ± 0.8 for the frequencies 1.189 kHz, 1.682 kHz, 2.379 kHz, 3.365 kHz, 4.759 kHz respectively, all post-hoc <.001 after FDR correction). There was also a significant effect of Group, with higher emissions for the Control group (p < .01) (see Figure 3.B).

*Distortion products Otoacoustic Emissions (DPOEA)* analysis revealed a main effect of Frequency (χ^2^(6) = 518.2, p < .001) depicted in supplementary Figure S2 (post-hoc with FDR correction). There was no effect of interaction with the factor Group.

Globally, we evidenced a deficit in the hearing abilities of participants with tinnitus compared with the controls. To compute a single score representing this group’s differences, we performed a Partial Least Squares Discriminant Analysis (PLS-DA), a supervised multivariate method that reduces dimensionality while selecting the most relevant variables to separate two groups. We included the auditory thresholds of the left and right ears from 125 Hz to 16 kHz. The first component explained 46% of the variance; the correlations between the component and the original variables are represented in Figure 3.C. As expected, the higher frequencies (above 2kHz) were more correlated with the component than the lower frequencies (below 2kHz), confirming that the hearing deficits between the groups is larger in higher pitches. We used the first component as a hearing loss score to control for the effect of this comorbidity in the linear models.

### Dysfunction of the arousal system associated with tinnitus

We used two complementary tasks (ANT and SART) to assess the attentional abilities of participants with and without tinnitus. All analysis included the emotional and hearing scores calculated in the preceding steps, to account for the potential effects of those tinnitus comorbidities on cognitive functioning.

#### Attentional Network task

Participants were very accurate in the task (mean accuracy ± standard error = 97.4% ± .03). Therefore, the analysis focused on the mean RTs, calculated after excluding error trials (incorrect and missing responses). The analyses of the *Endogenous orientation, Exogenous orientation and Flanker conflicts* revealed significant effects of the Cues (Wald χ^2^ = 195 - 1,670, *p* < .001), confirming that the experimental manipulations worked. Participants were faster with a valid cue (801 ± 9ms) than a double cue (845 ± 9ms), with a valid cue (795 ± 9ms) than after an invalid cue (882 ± 9ms), and with a congruent cue (748 ± 8ms) than an incongruent cue (913 ± 8ms). There were significant main Group effects (Wald χ^2^ = 4.9 - 5.9, *p* < .05), participants with tinnitus responding globally slowly than controls. There was no significant interaction between the Cues and the Group, indicating no evidence of dysfunction in these attentional subsystems associated with tinnitus.

In last, we analyzed the functioning of the arousal system: the alert and sustained attention. Analysis of the *Alerting* revealed significant effects of the Cue (Wald χ^2^ = 113.5, *p* < .001), of the Group (Wald χ^2^ = 4.6, *p* < .05) and a significant interaction between Group x Type of cue (Wald χ^2^ = 4.7, *p* < .05). As seen in Figure 4.A, Control participants responded faster than Tinnitus participants after a double cue (planned comparison: p < .05, Bonferroni correction), but there was no group difference in the absence of cue. This result indicates a lower speeding benefit of an alerting cue in presence of tinnitus.

**Figure 4:**
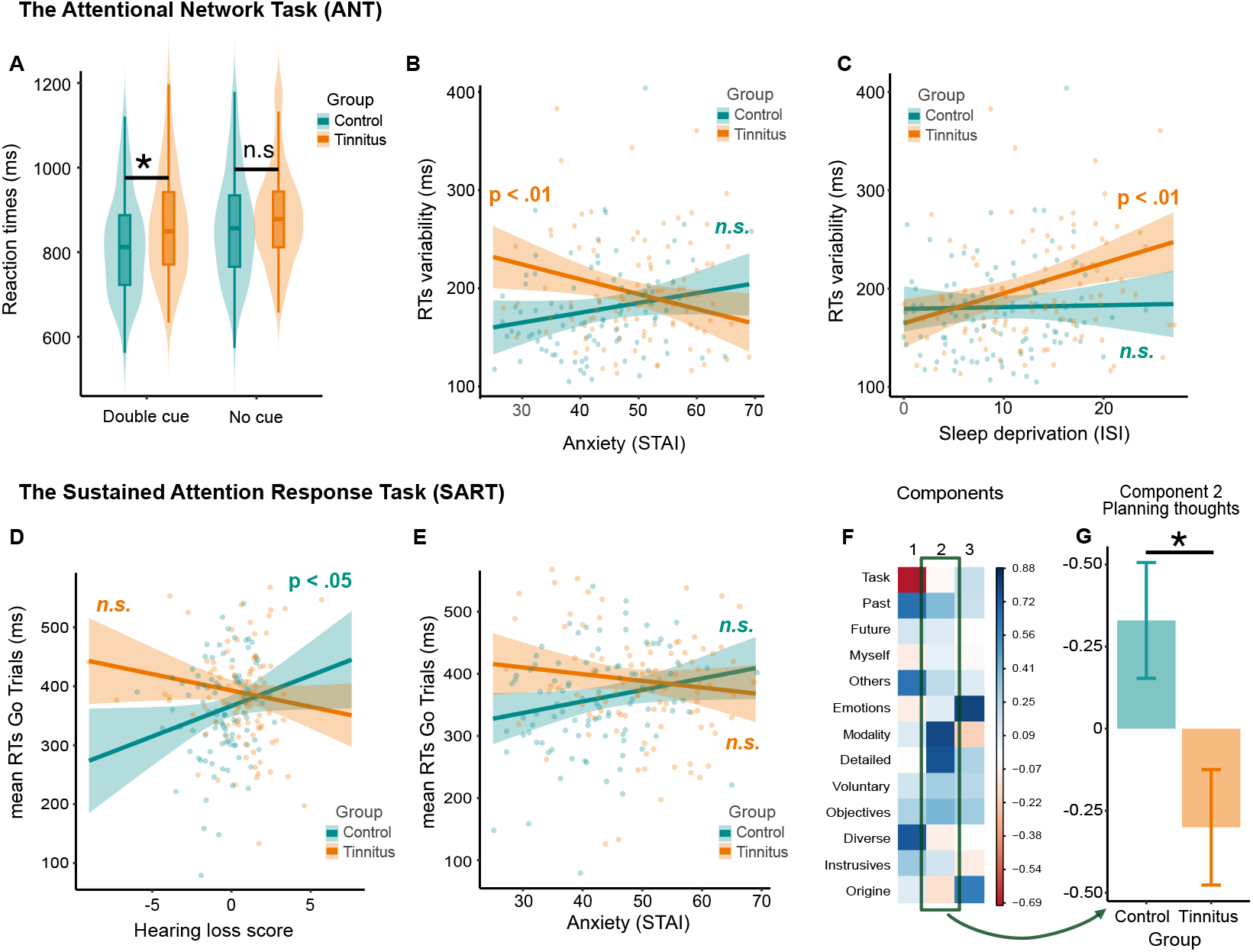
Attentional differences between tinnitus and control groups. A to C: ANT Results. **A**. Tinnitus group showed a reduced alerting effect compared to control, characterized by less sensitivity to double cues compared to a situation without cue. **B and C**. RTs variability, a measure of sustained attention, was higher in the tinnitus group. There was an interaction with both anxiety (B) and the sleep deprivation (C). Specifically, in tinnitus participants, sustained attention improved with increased anxiety and deteriorated with sleep deprivation, whereas control participants maintained stable performance across these variables. **D to G: Results of the SART. D and E**. Mean reaction times during the SART were significantly higher in the tinnitus group, indicating impaired sustained attention. Significant interactions were observed with hearing loss (D) and anxiety (E), demonstrating that these factors differently modulate of sustained attention between groups. **F**. Principal Component Analysis on thought content during SART task interruptions revealed three primary components: task-related thoughts (Component 1), planning-related thoughts (Component 2), and memory-related thoughts (Component 3). **G**. Between-group analysis of Component 2 revealed significantly more planning-related thoughts in the control group compared to the tinnitus group

The analysis of the *Sustained attention* revealed an effect of the Group (F(1,188) = 3.8, *p* = .05) with more variable RTs in the Tinnitus group. There was also an effect of Sleep disorder (F(1,188) = 6.3, *p* < .05), as well as an interaction between Group x Sleep disorder (F(1,188) = 4.6, *p* < .05): the RTs variability among participants with tinnitus increased with sleep disorders (slope different from zero: p<.01), which was not the case for control participants (Figure 4.B). Analysis also revealed an interaction between Group x Anxiety (F(1,188) = 7.9, p < .01): the RTs variability of participants with tinnitus decreased when anxiety increased (slope different from zero: p<.05), which was not the case for control participants (Figure 4.C). Globally, participants with tinnitus presented a deficit in sustained attention, modulated by their level of sleep and anxiety, which was not the case for control participants.

#### Sustained attention to Response task (SART)

We used the SART to study more closely the sustained attention abilities and evaluate mind wandering in this task thanks to series of probes.

*Sustained attention* is linked in this task to no go trials accuracy and Go trials RTs. The analysis of the *No Go trials Accuracy* revealed an effect of sleep (F(1,185) = 6.2, *p* < .05) and of hearing loss (F(1,185) = 4.9, *p* < .05), accuracy on No Go trials being higher with aging and hearing loss, and lower with sleep deprivation. There was no effect of the Group on this measure. Conversely, analysis of the *RTs Go trials* evidenced effects of the Group (F(1,185) = 4.37, p <.05) with slower RTs in the Tinnitus group, an interaction between Group and Anxiety (F(1,185) = 4.7, *p* < .05,) with RTs decreasing with anxiety in the tinnitus groups and increasing in the control group (slopes are not statistically different from 0) (Figure 4.D). Analysis also revealed an interaction between Group and Hearing loss (F(1,185) = 6.4, *p* < .05), with higher RTs associated with hearing loss in the control group only (slope p<.05), but not in the tinnitus group. (Figure 4.E).

The answers to the *Mind wandering* probe questions were first analyzed using a PCA, with the extraction of three components (see Figure 4.F). According to the highest loadings associated with each component, we can describe the first component as thoughts related to the task, to events from the past, thoughts on others and several subjects. The second component was thoughts using words, specific, on solution or objectives. The third was thoughts with an emotional valence, from information in their memory.

The analysis of the *first component* (task-related thoughts) *and third component* (memory-related thoughts) did not revealed effects of the Group. In opposition, the analysis of the *second component* (planning thoughts) indicated an effect of Group (F(1,185) = 5.4, p < .05), indicating that participants with higher education exhibited more planning-related thoughts, while the control group demonstrated significantly more planning thoughts than the tinnitus group (see Figure 4.G).

### Executive functions deficit is associated with hearing loss

Finally, we studied the executive functioning in presence of tinnitus, using the TMT and Stroop tests. Analysis of the *Trail making test (TMT)* and the *Stroop* revealed effect of Hearing loss (F(1,186) = 4.8-6.1, *p* = .05) with better performances associated with lower hearing loss. There was no effect of the factor Group.

### No evidence of links between the attentional scores and tinnitus characteristics

We tested if the scores that differ between groups (ANT Alerting effect, ANT RTs variability, and SART RTs) were linked to tinnitus characteristics, as its severity (measured with the THI score), its central pitch or duration (in years) by performing spearman correlation. We found no significant result.

## Discussion

In this study, we investigated the attentional functions (both endogenous and exogenous orientation), the arousal (alert and sustained attention) and the executive functions associated with tinnitus in a large sample size. As the tinnitus and control groups differed in hearing abilities, emotional state and sleep deprivation, those factors were controlled within the statistical model. We evidenced a deficit in phasic alertness, i.e. the speeding following the perception of a salient event, and a deficit in sustained attention in relation with anxiety, sleep and hearing loss. Unlike the data reported in the literature, we did not find a link between executive functions and tinnitus perception, but we did find an association with tinnitus-associated comorbidities—hearing loss and sleep deprivation—indicating that this cognitive deficit may be more closely related to these co-occurring conditions than to tinnitus itself.

Before addressing the main findings, it is important to consider the auditory and emotional profiles of participants with tinnitus in this study. Consistent with previous literature, individuals with tinnitus exhibited higher levels of stress, anxiety, depression, and sleep disturbances compared to controls (16). Despite pure-tone audiometry suggesting no hearing impairment in the tinnitus group (21), they showed subtle but consistent deficits in hearing function, including elevated thresholds and reduced distortion product amplitudes. Otoacoustic emissions remained comparable across groups. In the analysis of attention and executive tasks, we carefully accounted for these emotional, sleep-related, and auditory comorbidities. Because hearing loss in our sample predominantly affected high frequencies, we avoided using the conventional clinical hearing score (average thresholds from 500 Hz to 4 kHz), and instead employed a Partial Least Squares (PLS) approach to capture group differences more accurately. To further isolate the specific contribution of tinnitus, we included hyperacusis, a common comorbidity affecting approximately 80% of individuals with tinnitus (22) as a covariate. This comprehensive approach ensured that our conclusions were linked to the perception of tinnitus itself, rather than its associated hearing and emotional conditions.

Group differences in attentional functioning were mainly observable for alertness. Specifically, participants with tinnitus experienced reduced benefits from alerting cues, suggesting either a lower bump of phasic arousal following the warning cue or a diminished impact of this bump on performance. Moreover, participants with tinnitus exhibited slower reaction time in the SART and greater variability in the ANT, both of which are indicative of diminished sustained attention (23, 24). These results suggest that tinnitus negatively impacts tonic alertness, maybe reflecting a poorer internal ability to modulate arousal to an optimal level for task performance. Interestingly, anxiety and sleep deprivation effects on sustained attention, while expected, were significantly more pronounced in the tinnitus group than in the control group. This finding suggests that individuals with tinnitus are less capable than controls to compensate or to cope with the detrimental effects of sleep deprivation and anxiety on performance.

Patterns of mind-wandering also differed between groups. Control participants reported more frequent problem-solving thoughts during the task, while those with tinnitus were less likely to engage in such task-unrelated thoughts. This suggests that, despite similar accuracy rates, individuals with tinnitus may have had fewer cognitive resources at their disposal, making it more difficult to shift attention away from the task compared to control participants. This is also in line with a potential disruption of tonic arousal in tinnitus, as arousal mediates to amount of available cognitive capacity during a task (25).

Phasic alertness and sustained attention are both underpinned by a common neurobiological mechanism: the locus coeruleus–norepinephrine (LC-NE) system (7, 27). This subcortical system critically regulates cortical excitability and sensory processing through distinct activity patterns: brief phasic bursts triggered by salient events generate momentary alerting responses, while sustained tonic activity maintains general arousal and vigilance, thereby supporting sustained attention. LC-NE effects on cognition follows an inverted-U curve (28). Moderate LC-NE activity enhances attention to an optimal level, whereas low activity (e.g., due to fatigue or drowsiness) can lead to mind-wandering and attentional lapses. Excessive levels (e.g., under stress) produce hypervigilance and distractibility (Figure 5). Our study revealed that tinnitus patients exhibited both reduced sustained attention, interacting with sleep deprivation and anxiety, and decreased phasic alertness. These findings collectively suggest LC-NE system dysregulation in tinnitus: a chronic hypoactivity within this critical neuro-modulatory system could explain why sleep deprivation and anxiety exhibit compounded effects in tinnitus compared to the general population.

**Figure 5:**
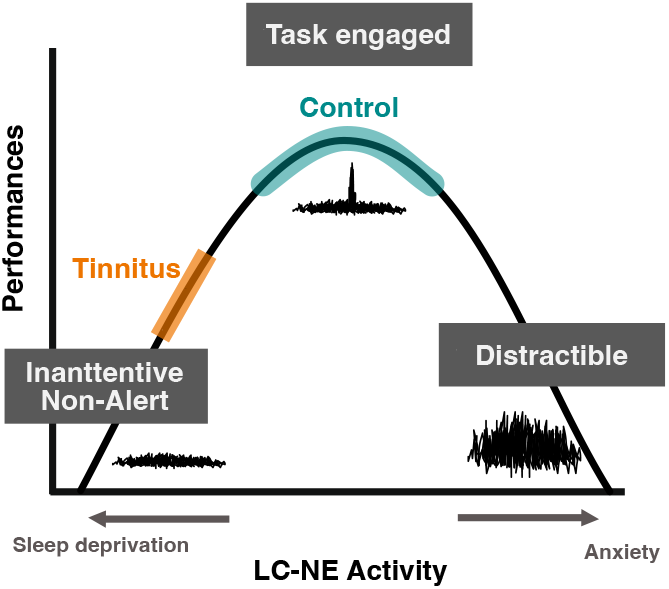
Activity of the LC-NE and attention. Alertness (phasic arousal) and sustained attention (tonic arousal) are related to the activity of the LC-NE. Their level impact cognitive performances with an inverted U-shape relationship. Performances are poor at low and high levels of LC. Based on our results, we hypothesized that individuals with tinnitus have a lower level of LC compared to controls.

Executive function deficits have been repeatedly reported in individuals with tinnitus (13, 14). However, the evidential strength of these findings remains limited, largely due to inadequate control of comorbidities and small sample sizes. Contrary to previous reports, we found no direct link between tinnitus and executive dysfunction; rather, deficits in inhibition and shifting were attributed to hearing loss and sleep deprivation. This is consistent with studies showing associations between hearing loss and executive function both in the general population (29, 30) and in tinnitus samples (31). Our findings suggest that previously observed executive deficits may reflect the impact of comorbidities rather than tinnitus itself, underscoring the importance of rigorous comorbidity control in cognitive studies of tinnitus.

In this study, we identified an association between tinnitus perception and dysfunctions in the arousal system, in interaction with sleep and anxiety. A remaining question is to account causality links between those symptoms. The classical explanation is that tinnitus disrupts attentional processes by constantly capturing attention trough high saliency, subsequently leading to sleep deprivation and anxiety (16, 32). Conversely, pre-existing attentional dysfunction might contribute to chronic tinnitus development (33, 34). Sedley and collaborators propose that attention difficulties can lead to an abnormal focus on tinnitus, increase its salience, alter perceptual priors and facilitate chronicity. A third possibility is that tinnitus perception and the dysregulation of attention, sleep and anxiety share a common underlying cause. Our results implicate the arousal system in tinnitus pathophysiology, a system intimately connected with attention, sleep regulation and anxiety. We hypothesize that a dysregulation of this system could simultaneously precipitate all three conditions. Further studies, particularly longitudinal designs, are needed to clarify the directionality and causal relationships among tinnitus, attention, sleep, and anxiety.

By rigorously controlling for key comorbidities, namely hearing loss, emotional distress, and sleep disturbances thanks to a large sample size, our study succeeded in disentangling the cognitive deficits genuinely associated with tinnitus from those attributable to its frequent comorbid conditions. We propose a novel theoretical framework in which dysregulation of arousal, likely supported by the locus coeruleus– norepinephrine (LC-NE) system, may mediate both tinnitus perception and its interactions with sleep and anxiety. This integrated model opens new avenues for understanding and targeting the cognitive and emotional burden of tinnitus, that deserve to be further investigated.

## Materials and Methods

### Participants

Two hundred participants recruited at the Hôpital Européen Georges Pompidou (HEGP, Paris) were divided in two groups: a Tinnitus group (n = 100), with individuals who have experienced subjective (non-pulsatile) chronic tinnitus for more than three months, and a Control group (n = 100), comprising individuals without tinnitus matched to the tinnitus group in terms of age and education level. Inclusion criteria were as follows: age between 18 and 65 years, absence of severe or profound hearing loss (no unilateral mean hearing loss exceeding 65 dB at 0.5 kHz, 1 kHz, 2 kHz, and 4 kHz (21)), normal or corrected-to-normal vision, absence of neurological disorders, and fluency in French. The study was approved by an institutional review board (CPP Sud-Est VI, Clermont-Ferrand, France; reference: AU 1643) and registered on ClinicalTrials.gov (first posted on 15/01/2021, identifier NCT04717388). All procedures were conducted in accordance with relevant guidelines and regulations. Informed written consent was obtained from all participants prior to participation.

### Socio-emotional and demographics

Information about participants’ demographic (age, sex, educational level), mental health with the Handicap Anxiety and Depression State (HADS) (35), the Spielberger State–Trait Anxiety Inventory (STAI) (36), the Perceived Stress scale (PSS-4) (37) and sleep quality with the Insomnia Severity Scale (ISI) (38) were collected.

### Auditory and tinnitus evaluations

All participants underwent an auditory evaluation, including a pure tone audiometry testing frequency from 125Hz, 250Hz, 500Hz, 750 Hz, 1kHz, 1.5kHz, 2kHz, 3kHz, 4kHz, 6kHz, 8kHz, 12kHz and 16kHz using the MedRx AVANT software. Auditory thresholds at the high frequencies (12kHz and 16kHz) were measured in dB HL and converted in SL using norms (39). Participants who did not perceived high frequencies were attributed the maximum value. Participants also performed oto-emissions at 1.189 kHz, 1.682 kHz, 2.379 kHz, 3.365 kHz, 4.759 kHz, and distortion products at 1.230 kHz, 1.641 kHz, 2.461 kHz, 3.281 kHz, 4.915 kHz, 6.555 kHz, 8.196 kHz. Measures were collected using the Interacoustics Eclipse with the OAE-Suite software.

Hyperacusis was evaluated using the Khalfa questionnaire (40) and a measure of the discomfort level at 1kHz in each ear. Patients with tinnitus answered questions about the tinnitus characteristics (i.e. time since onset, localization, type of sound…) taken from the ESIT-SQ (41), filled the Tinnitus Handicap Inventory questionnaire (THI) (42). Additionally, they performed a tinnitus matching that allows to create a sound matching the tinnitus percept by choosing the loudness, the central pitch, a type of noise around this central pitch and a rhythm, using the MedRx tinnometer.

### Attentional Network task

We used the revised Attentional Network task (ANT-R) (8). The task presents stimuli consisting of five horizontal black arrows in a row (one central arrow with four flankers) pointing leftward or rightward. The task is to identify as quickly and accurately as possible the central arrow’s direction by pressing a key with the index finger (left hand for a leftward direction, right hand for a rightward direction). Reaction times (RTs) were collected up to 1700ms after target onset.

A cue in the form of a 100ms flashing box could precede the target (by 0, 400, or 800ms) under three conditions: i) No-cue: no flashing cue box. ii) Double-cue: both boxes flash, providing temporal information only, iii) Spatial-cue: one box flashes, offering both temporal and spatial information. Intertrial intervals ranged from 2000-12000ms (mean=4000ms). A fixation cross was visible at the center of the screen through-out the duration of the task. The protocol is illustrated in Figure 1.

The experimental conditions include the factors: Cue (no-cue, double-cue, invalid spatial-cue, valid spatial-cue, with three time more presentation of valid-spatial cue than the other combination), Target location (left, right), Target directions (left, right), Flanker congruencies (congruent, incongruent) and the Cue-to-target delays (0, 400, and 800ms). The combination of those conditions leads to 144 trials, split into 2 runs of 72 counterbalanced trials. Based on those conditions, we defined different conditions of interest. The **valid/invalid trials** were trials with spatial cues (left/right), with a similar/different spatial position than the target (left/right). The **congruent/incongruent trials** were trials with similar/different orientation of the target arrow (left/right) and the flanker arrows (left/right). Different scores were computed based on the different conditions to assess the function of each attentional subcomponents.

- **Endogenous orientation:** Speeding benefit of a lateralized valid cue, calculated as: *RTs_double cue - RTs_valid cue*.
- **Exogenous orientation**: Speeding benefit of a valid cue compared to an invalid one, calculated as: *RTs_invalid cue - RTs_valid cue*.
- **Alerting Effect**: Speeding benefit of an alerting cue calculated as: *RTs_no cue – RTs_double cue*.
- **Sustained Attention**: Variability in reaction times, calculated as *standard deviation of RTs* throughout the task.
- **Executive Control (Flanker Conflict Effect)**: Cost of inhibiting the distracting incongruent flanker**s**, calculated as: *RT_flanker incongruent – RT_flanker congruent*

### Sustained attention to Response task (SART)

We used the ordered SART, which is a Go/No-Go paradigm (Figure 2.A). This task is composed of sequential presentation of numbers from 1 to 9 were presented sequentially. Participants had to press a button whenever a number appears, except when the number is three, in which case they must withhold their response. Each number remained on the screen for one second, followed by a one-second display of a circle. Each number was presented 33 times throughout the task. Periodically, the task paused, and participants responded to a series of 13 questions adapted from (43) assessing mind wandering (Figure 2.B). Participants had to rate on a 1 to 10 scale whether they were their thoughts were: *on the task, about past events, about future events, about themselves, about others, negative or positive, in form of images or words, detailed and specific, about solutions to problems, about one or several subjects, intrusive, linked to information from the environment or their memory*, and finally *if they went into their thoughts voluntarily*. There were 10 sets of questions. We calculated the accuracy on the no-go trials, the mean RTs as indicators of participants sustained attention. Answers to the probe questions were analyzed with a principal component analysis, from which we extracted three components in line with (44).

### Executive tasks: the Stroop and Trail Making Test (TMT)

We used the classical Stroop color-word test to evaluate inhibition (44). The task consists of three main conditions: 1. Color condition – identify the color of rectangles. 2. Reading condition – reading color words printed in black ink. 3. Interference Condition –naming the color of the ink of color words while inhibiting the reading of the word (e.g., if the word “RED” is written in blue ink, the answer is “blue”). Participants were instructed to be as quick and accurate as possible. The time difference between the reading and interference conditions represented the inhibition ability.

The TMT is composed of two conditions assessing cognitive flexibility (46). In condition A, participants had to connect numbers in ascending order as fast as possible (1, 2, 3…). In condition B, participants had to alternate between numbers and letters in ascending and alphabetical order, respectively (1, A, 2, B…). Participants were instructed to do the task as quickly and accurately as possible. The time difference between the two conditions reflects cognitive flexibility ability.

### Statistical analysis

The hearing thresholds, oto-emissions and distortion products were analyzed using linear mixed-effects model including as fixed effects the factors Age, Sex (Female/Male), Ear (Left/Right), Group (Tinnitus/Control), Frequency, the interactions between the factors Frequency × Group, Age × Group, and the factor Subject as random effect.

Data were analyzed using a linear mixed-effects model (LMM), including as fixed effects the factors Age, Sex, Education level, Group, Hearing loss, Hyperacusis, Sleep disorder, Anxiety, Cue and the interactions between the factors Cue x Group. The factor Subject was included as a random factor. We applied a logarithmic transformation on the means RTs to meet the LMM assumptions.

All analysis except the sustained attention included a two levels factor cue, with levels for each attentional subfunction: valid / invalid for Exogenous orientation, valid / no cue for Endogenous orientation, no cue/double cue for the Alerting and Conflict / no conflict for the Executive.

In the SART, the TMT and Stroop, all data were analyzed using linear model with the factors Age, Sex, Education level, Group, Hearing loss, Hyperacusis, Sleep disorder, Anxiety, and the interactions between the Group and the factors Hearing loss, Sleep disorder and Anxiety We applied a logarithmic transformation on the means RTs to meet the model assumptions.

## Acknowledgments

The project was supported by the Fondation pour l’Audition (FPA RD-2019-10). This work has benefited from a French government grant managed by the Agence Nationale de la Recherche under the France 2030 program, reference ANR-23-IAHU-0003. LH was funded by the European Union’s Horizon Europe Framework Programme (HORIZON) under the Marie Skłodowska-Curie Postdoctoral Fellowship (grant No. 101146406), and by the Fondation des Gueules Cassées.

## Author Contributions

All authors conceived and designed the study. LH collected the data, performed the statistical analyses, and drafted the manuscript. All authors were involved in the interpretation of data, and critical revision of the manuscript.

## Figures and Tables

## Supplementary Materials

**Figure S1:**
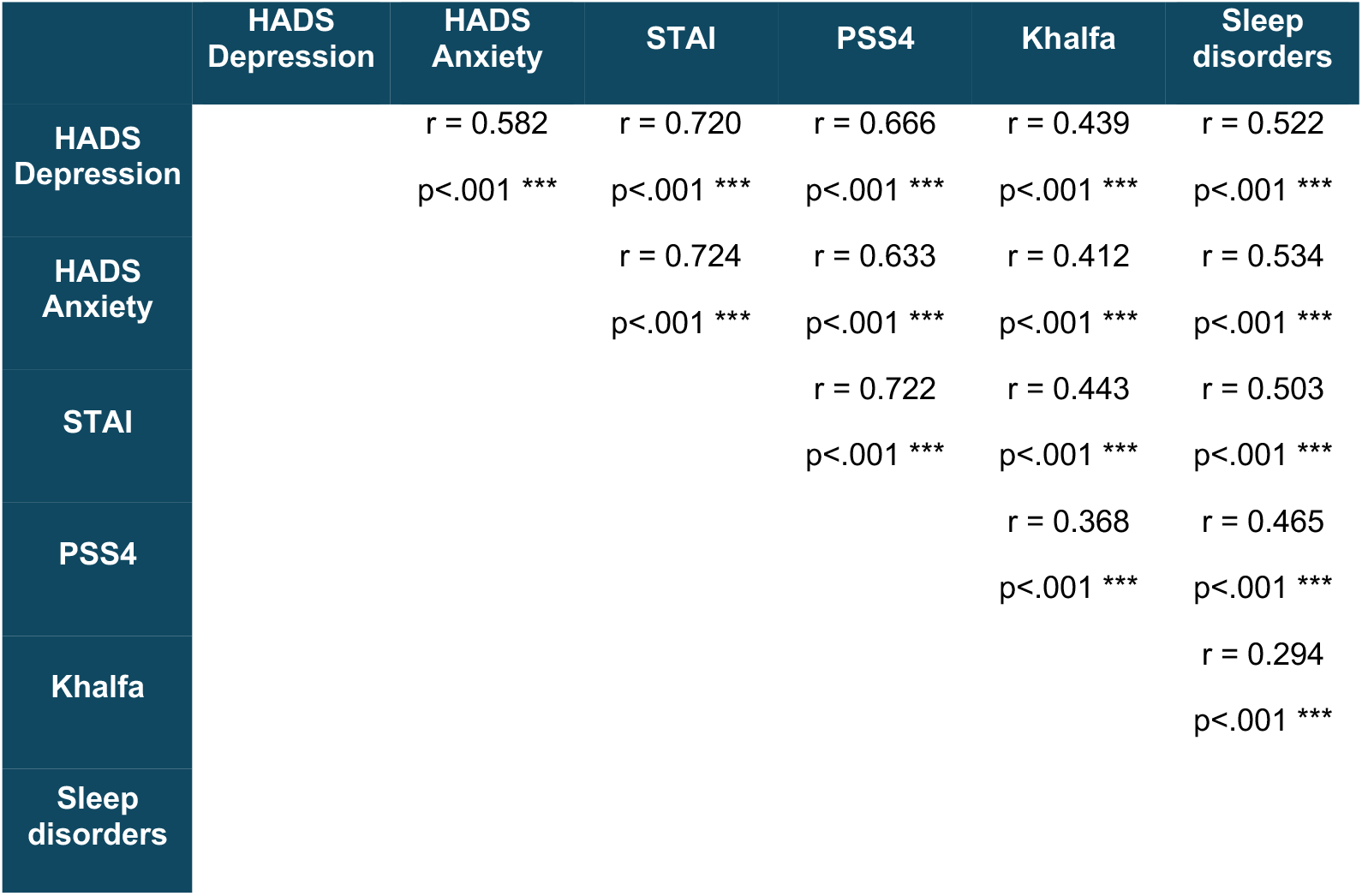
Emotional evaluation: matrix of correlation between the difference measures

**Figure S2:**
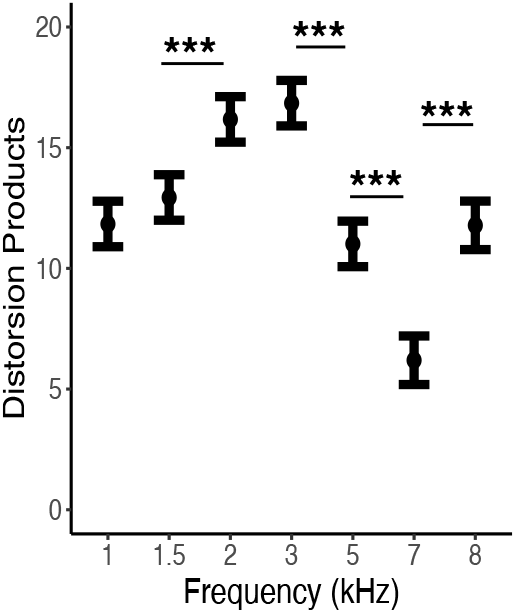
Distortion products Otoacoustic Emissions (DPOEA) level of each tested frequency

## References

1. C. M. Jarach, et al., Global Prevalence and Incidence of Tinnitus: A Systematic Review and Meta-analysis. JAMA Neurol 79, 888 (2022).

2. D. Baguley, D. McFerran, D. Hall, Tinnitus. The Lancet 382, 1600–1607 (2013).

3. D. A. Hall, et al., A narrative synthesis of research evidence for tinnitus-related complaints as reported by patients and their significant others. Health Qual Life Outcomes 16, 61 (2018).

4. C. A. Bauer, Tinnitus. New England Journal of Medicine 378, 1224–1231 (2018).

5. K. J. Trevis, N. M. McLachlan, S. J. Wilson, Cognitive mechanisms in chronic tinnitus: Psychological markers of a failure to switch attention. Frontiers in Psychology 7 (2016).

6. M. D. Rosenberg, E. S. Finn, D. Scheinost, R. T. Constable, M. M. Chun, Characterizing Attention with Predictive Network Models. Trends in Cognitive Sciences 21, 290–302 (2017).

7. S. J. Sara, S. Bouret, Orienting and Reorienting: The Locus Coeruleus Mediates Cognition through Arousal. Neuron 76, 130–141 (2012).

8. J. Fan, et al., Testing the behavioral interaction and integration of attentional networks. Brain and Cognition 70, 209–220 (2009).

9. R. S. Hallam, L. Mckenna, L. Shurlock, Tinnitus impairs cognitive efficiency. International Journal of Audiology 43, 218–226 (2004).

10. C. Stevens, G. Walker, M. Boyer, M. Gallagher, Severe tinnitus and its effect on selective and divided attention: Acufeno severo y sus efectos sobre la atención selectiva y dividida. International Journal of Audiology 46, 208–216 (2007).

11. G. Andersson, A. Khakpoor, L. Lyttkens, Masking of tinnitus and mental activity. Clin Otolaryngol 27, 270–274 (2002).

12. A. Heeren, et al., Tinnitus specifically alters the top-down executive control sub-component of attention: Evidence from the Attention Network Task. Behavioural Brain Research 269, 147–154 (2014).

13. S. Tegg-Quinn, R. J. Bennett, R. H. Eikelboom, D. M. Baguley, The impact of tinnitus upon cognition in adults: A systematic review. International Journal of Audiology 55, 533–540 (2016).

14. N. A. Clarke, H. Henshaw, M. A. Akeroyd, B. Adams, D. J. Hoare, Associations Between Subjective Tinnitus and Cognitive Performance: Systematic Review and Meta-Analyses. Trends in Hearing 24, 233121652091841 (2020).

15. N. Mohamad, D. J. Hoare, D. A. Hall, The consequences of tinnitus and tinnitus severity on cognition: A review of the behavioural evidence. Hearing Research 332, 199–209 (2016).

16. K. J. Trevis, N. M. Mclachlan, S. J. Wilson, A systematic review and meta-analysis of psychological functioning in chronic tinnitus. Clinical Psychology Review 60, 62–86 (2018).

17. H. Vasudevan, K. Ganapathy, H. P. Palaniswamy, G. Searchfield, B. Rajashekhar, Systematic review and meta-analysis on the effect of continuous subjective tinnitus on attention and habituation. PeerJ 9, e12340 (2021).

18. L. Hobeika, et al., Tinnitus risk factors and its evolution over time. Nat Commun 16, 4244 (2025).

19. T. D. Griffiths, et al., How Can Hearing Loss Cause Dementia? Neuron 108, 401–412 (2020).

20. I. H. Robertson, T. Manly, J. Andrade, B. T. Baddeley, J. Yiend, ‘Oops!’: Performance correlates of everyday attentional failures in traumatic brain injured and normal subjects. Neuropsychologia 35, 747–758 (1997).

21. B. O. Olusanya, A. C. Davis, H. J. Hoffman, Hearing loss grades and the International classification of functioning, disability and health. Bull. World Health Organ. 97, 725–728 (2019).

22. C. Cederroth, et al., Association between Hyperacusis and Tinnitus. JCM 9, 2412 (2020).

23. D. Smilek, J. S. A. Carriere, J. A. Cheyne, Failures of sustained attention in life, lab, and brain: Ecological validity of the SART. Neuropsychologia 48, 2564–2570 (2010).

24. A. Yamashita, et al., Variable rather than extreme slow reaction times distinguish brain states during sustained attention. Sci Rep 11, 14883 (2021).

25. D. Kahneman, Attention and effort (Prentice-Hall, Inc, 1973).

26. S. L. Leong, et al., The potential interruptive effect of tinnitus-related distress on attention. Sci Rep 10, 11911 (2020).

27. W. Sturm, K. Willmes, On the Functional Neuroanatomy of Intrinsic and Phasic Alertness. NeuroImage 14, S76–S84 (2001).

28. G. Aston-Jones, J. D. Cohen, An integrative theory of locus coeruleus-norepinephrine function: adaptive fain and optimal performance. Annu. Rev. Neurosci. 28, 403–450 (2005).

29. S. A. M. Valentijn, et al., Change in Sensory Functioning Predicts Change in Cognitive Functioning: Results from a 6-Year Follow-Up in the Maastricht Aging Study. J American Geriatrics Society 53, 374–380 (2005).

30. F. R. Lin, et al., Hearing loss and cognition in the Baltimore Longitudinal Study of Aging. Neuropsychology 25, 763–770 (2011).

31. A. M. Pajor, E. A. Ormezowska, M. Jozefowicz-Korczynska, The impact of co-morbid factors on the psychological outcome of tinnitus patients. Eur Arch Otorhinolaryngol 270, 881–888 (2013).

32. H. Gu, W. Kong, H. Yin, Y. Zheng, Prevalence of sleep impairment in patients with tinnitus: a systematic review and single-arm meta-analysis. Eur Arch Otorhinolaryngol 279, 2211–2221 (2022).

33. L. E. Roberts, F. T. Husain, J. J. Eggermont, Role of attention in the generation and modulation of tinnitus. Neuroscience & Biobehavioral Reviews 37, 1754–1773 (2013).

34. W. Sedley, K. J. Friston, P. E. Gander, S. Kumar, T. D. Griffiths, An Integrative Tinnitus Model Based on Sensory Precision. Trends in Neurosciences 39, 799–812 (2016).

35. I. Bjelland, A. A. Dahl, T. T. Haug, D. Neckelmann, The validity of the Hospital Anxiety and Depression Scale An updated literature review. Journal of Psychosomatic Research (2002).

36. T. M. Marteau, H. Bekker, The development of a six-item short-form of the state scale of the Spielberger State—Trait Anxiety Inventory (STAI). British J Clinic Psychol 31, 301–306 (1992).

37. S. Cohen, T. Kamarck, R. Mermelstein, A Global Measure of Perceived Stress. Journal of Health and Social Behavior 24, 385 (1983).

38. C. M. Morin, G. Belleville, L. Bélanger, H. Ivers, The Insomnia Severity Index: Psychometric Indicators to Detect Insomnia Cases and Evaluate Treatment Response. Sleep 34, 601–608 (2011).

39. A. Rodríguez Valiente, A. Trinidad, J. R. García Berrocal, C. Górriz, R. Ramírez Camacho, Extended high-frequency (9–20 kHz) audiometry reference thresholds in 645 healthy subjects. International Journal of Audiology 53, 531–545 (2014).

40. S. Khalfa, et al., Psychometric Normalization of a Hyperacusis Questionnaire. ORL 64, 436–442 (2002).

41. E. Genitsaridi, et al., Standardised profiling for tinnitus research: The European School for Interdisciplinary Tinnitus Research Screening Questionnaire (ESIT-SQ). Hearing Research 377, 353– 359 (2019).

42. W. Newman, G. P. Jacobson, J. B. Spitzer, Development of the Tinnitus Handicap Inventory. Arch Otolaryngol Head Neck Surg 122, 143–148 (1996).

43. D. Konu, et al., A role for the ventromedial prefrontal cortex in self-generated episodic social cognition. NeuroImage 218, 116977 (2020).

44. J. Smallwood, et al., Representing Representation: Integration between the Temporal Lobe and the Posterior Cingulate Influences the Content and Form of Spontaneous Thought. PLoS ONE 11, e0152272 (2016).

45. J. R. Stroop, Studies of interference in serial verbal reactions. Journal of Experimental Psychology 18, 643–662 (1935).

46. R. M. Reitan, Validity of the Trail Making Test as an Indicator of Organic Brain Damage. Percept Mot Skills 8, 271–276 (1958).

